# Fluorescence *in situ* hybridization reveals endophytic and epiphytic root colonization of the novel plant growth-promoting bacterium *Citrobacter sedlakii* CESi7

**DOI:** 10.64898/2026.06.28.735065

**Authors:** Hiromu Inoue, Manami Maeda, Tomonori Koga, Zahra Salman, Clament Fui Seung Chin, Mohd. Huzairi Mohd. Zainudin, Norhayati Bt. Ramli, Mohd Ali Hassan, Yukihiro Tashiro, Kenji Sakai

## Abstract

Plant growth-promoting bacteria are gaining significant attention as promising biofertilizers. However, the inconsistency between *in vitro* plant growth-promoting traits and actual field performance remains a challenge, driven partly by a limited understanding of *in situ* colonization. This study characterized the colonization patterns of *Citrobacter sedlakii* CESi7, a novel plant growth-promoting bacterium, isolated from oil palm waste compost, during *Brassica rapa* cultivation. The *in situ* behavior of CESi7 was observed in both sterilized medium and non-sterilized soil using fluorescence *in situ* hybridization with a strain-targeting probe. The results revealed that CESi7 can establish both epiphytic and endophytic populations that transiently colonize roots. In a sterilized medium, CESi7 was widely distributed throughout the root tissues. Conversely, in non-sterilized soil, the bacterium formed dense aggregates specifically at the root tips. This study provides direct microscopic evidence of the colonization strategy of CESi7, offering crucial insights for its development as an effective biofertilizer.

---

Plant growth-promoting bacteria (PGPB) inhabit various plant parts, including the rhizosphere soil, root tissues, seeds, and leaves, thereby enhancing plant growth (Bashan *et al*. 1998). Important plant growth-promoting traits can be classified into two mechanistic categories: direct functions, such as mineral solubilization, nitrogen fixation, and phytohormone production; and indirect functions, such as the production of siderophores and antibiotics (Gupta *et al*. 2015). Consequently, PGPB represent promising biofertilizers for advancing sustainable and environmentally friendly agricultural practices (Gupta *et al*. 2015). To date, PGPB have been tested on various crops, including wheat (Gontia-Mishra *et al*. 2016), oats (Sapre *et al*. 2018), and rice (Nguyen *et al*. 2003).

The plant rhizosphere is enriched with diverse microorganisms such as bacteria, and serves as a key site for plant–microbe interactions (Bais *et al*., 2006; Reinhold-Hurek *et al*. 2015). Rhizosphere colonization by PGPB is crucial for plant growth promotion for several reasons: (i) beneficial substances produced by PGPB are directly supplied to the host plant without being consumed or removed by other microorganisms; (ii) PGPB securely attached to roots are less susceptible to be displaced by water, thereby ensuring stable plant growth promotion; and (iii) the occupation of the rhizosphere niche by antagonistic PGPB decreases the risk of colonization by pathogenic microorganisms (Mia *et al*. 2010). Therefore, clarifying the spatiotemporal dynamics of PGPB colonization in the rhizosphere using microscopic observations provides a fundamental basis for agricultural applications.

In our previous study, *Citrobacter sedlakii* CESi7, a novel PGPB isolated from oil palm waste compost, demonstrated a high capacity for nitrogen fixation, phosphorus and silicate solubilization, as well as indole-3-acetic acid and siderophore production *in vitro* (Chin *et al*. 2017). Furthermore, inoculation with strain CESi7 significantly increased the growth of *Brassica rapa* (komatsuna) in a pot cultivation test (Chin *et al*. 2017). To facilitate the practical application of strain CESi7 as a biofertilizer, a comprehensive understanding of its colonization dynamics within root systems and surrounding soil is essential. In this study, we developed a fluorescence *in situ* hybridization (FISH) probe targeting strain CESi7 to visualize and quantify its *in situ* behavior in both sterilized and non-sterilized environments.

## MATERIALS AND METHODS

### Bacterial strains and culture conditions

*C. sedlakii* CESi7 was grown in trypticase soy broth (Difco, Becton Dickinson, Franklin Lakes, NJ, USA) adjusted to an initial pH of 7.3 on an orbital shaker (230 rpm) at 37 °C for 24 h. *Rheinheimera tangshanensis* JCM14254^T^ was cultivated at 28 °C and an initial pH of 7.0 for 18 h in Luria-Bertani broth containing 10 g/L tryptone (Nacalai Tesque, Kyoto, Japan), 5 g/L yeast extract (Oriental Yeast, Tokyo, Japan), and 10 g/L NaCl. *Mixta theicola* NBRC110557^T^ was cultivated at 35 °C and an initial pH of 7.0 for 24 h in No. 802 medium containing 10 g/L hipolypeptone (Nihon Pharmaceutical, Tokyo, Japan), 2 g/L yeast extract (Oriental Yeast), and 1 g/L MgSO_4_·7H_2_O. *Escherichia coli* DH5α was cultured at 37 °C and an initial pH of 7.0 for 24 h in nutrient broth containing 10 g/L peptone (Difco), 10 g/L beef extract (Nacalai Tesque), and 5 g/L NaCl.

### Probe design for FISH

The 16S rRNA gene sequence of strain CESi7 was submitted to the EzBioCloud Database (https://www.ezbiocloud.net/identify) to retrieve 16S rRNA gene sequences from the top 10 most closely related strains. Subsequently, these sequences were subjected to multiple sequence alignment using the ClustalW algorithm implemented in MEGA X (https://www.megasoftware.net). Partial 35-mer regions exhibiting a high number of mismatches between strain CESi7 and the non-target type strains were selected as candidate regions. From these, 18-mer probe candidate regions were selected by sliding every 1 mer across the 35-mer sequences. These candidates were then evaluated in terms of mishit rate, brightness class (Fuchs *et al*. 1998), and melting temperature (*T*_m_). The mishit rate was calculated using the Probe Match application of the Ribosomal Database Project (https://rdp.cme.msu.edu/probematch/search.jsp). The candidate region was determined with a low mishit ratio, brightness classes IV or higher, and a *T*_m_ of ca. 60 °C. Finally, the 3 mer of the flanking region of the target sequence was extended from the 5′ end, yielding a probe targeting strain CESi7 (CITR906) (Table 1). Probes EUB338 (Amann *et al*. 1990) and NONEUB (Wallner *et al*. 1993) labeled with rhodamine red X (ROX) were used as universal probes for bacteria and negative controls, respectively (Table 1). The probes were purchased from Hokkaido System Science (Sapporo, Japan).

**Table 1.**
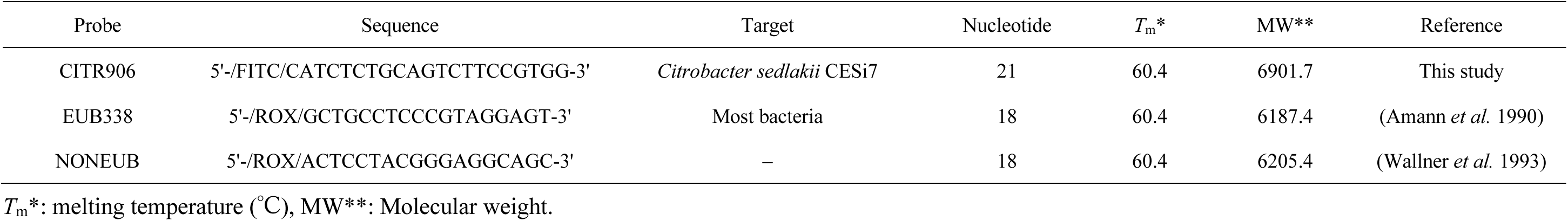
Probes used in this study.

### Plant cultivation conditions

*B. rapa* was cultivated in either sterilized medium or non-sterilized soil under greenhouse conditions (25 °C and 70% relative humidity). The experiment was conducted using a completely randomized design as described previously (Chin *et al*. 2017). Murashige and Skoog (MS) medium containing 0.43% (w/v) MS basal salt mixture (Merck KGaA, Darmstadt, Germany) and 0.8% (w/v) agar at an initial pH of 6.0 was used (Khan *et al*. 1999). After autoclaving at 110 °C for 20 min, 150 mL of the medium was transferred to ca. 950-mL sterile vessels (PHYTOCOM-32; PhytoTechnology Laboratories, KS, USA) and completely solidified. The vessels were covered with sterile ventilated lids (Closure PHYTOCON; PhytoTechnology Laboratories). For the non-sterilized soil, commercial black soil (Gou building material, Utsunomiya, Japan) was used after sieving through a 2-mm stainless steel sieve. The cultivation pots (inner diameter, 13 cm) were filled with 500 g of black soil per pot.

### Seed sterilization and inoculation

Viable seeds were selected using a float test. The seeds that sank were collected and sterilized with 1% (w/v) sodium hypochlorite for 30 min, followed by 70% (v/v) ethanol for 5 min. After washing five times with sterile distilled water (ddH_2_O), 10 seeds were transferred to a 2.0-mL centrifuge tube. One milliliter of strain CESi7 cell suspension in phosphate-buffer saline (PBS) buffer (8 g/L NaCl, 0.2 g/L KCl, 1.44 g/L Na_2_HPO_4_, and 0.24 g/L KH_2_PO_4_; pH 7.4), adjusted to 3.0 × 10^8^ cells/mL, was added to the tubes and incubated at 37 °C for 1 h. After the suspension was removed by pipette, the cell-coated seeds were dried overnight at 37 °C. The inoculated seeds were then sown in the sterilized medium (5 seeds/pot) or non-sterilized soil (10 seeds/pot). During cultivation in soil, ddH_2_O was supplied daily to maintain the soil water content at 70% (w/w) saturation.

### Sampling of roots and soil

The roots, rhizosphere soil, and bulk soil were collected after one, three, and five weeks of cultivation. The roots were gently washed with ddH_2_O and divided into two sections: the root tips (apical 5 mm) and the remaining upper portions, referred to as the mature zones. Rhizosphere soil was collected within 1 cm of the root surface, whereas bulk soil was collected from the remaining area. The soil samples were collected from a depth of 3 cm below the soil surface. Only the roots were collected after one, three, and five weeks of cultivation in the medium, as described above.

### Fixation of cells, roots, and soil

For pure culture samples, the cells from 120 mL of culture broth were fixed in 4% (w/v) paraformaldehyde (Wako, Osaka, Japan). Following incubation at 4 °C for 4 h, cells were collected via centrifugation at 12,000 × *g* for 5 min and washed twice with 50 mL of PBS buffer (pH 7.4). The fixed cells were stored in a 1:1 mixture of 99% (v/v) ethanol and PBS buffer (pH 7.4) at −20 °C until use.

Collected roots and soil subsamples (0.1 g each) were fixed with 4% (w/v) paraformaldehyde (Wako) at 4 °C for 8 and 12 h, respectively. The fixed roots and soil were washed twice with 750 μL and 1 mL of PBS buffer (pH 7.4), followed by centrifugation at 12,000 × *g* for 5 min. The fixed samples were stored in a 1:1 mixture of 99% (v/v) ethanol and PBS buffer (pH 7.4) at −20 °C until analysis.

### FISH

The fixed roots were immersed in a series of sucrose solutions (10% and 20% [w/v]) for 1 h each and subsequently embedded in O.C.T. compound (Sakura Finetek Japan, Tokyo, Japan). The embedded samples were frozen and cut into 20-µm thick sections using a CryoStar NX70 (Thermo Fisher Scientific, MA, USA). The root sections were mounted on MAS-coated glass slides (Matsunami, Osaka, Japan) and stored at 4 °C until use. For fixed cell cultures, rhizosphere and bulk soil, 4-μL aliquots of 10-fold diluted suspensions in PBS buffer (pH 7.4) were applied to 12-well glass slides (Matsunami). The slides were then dried at 46 °C for 20 min to ensure sample adherence.

For cell-wall permeabilization, samples were treated with lysozyme solution (20 mg/mL lysozyme, 100 mM Tris-HCl, and 50 mM EDTA; pH 8.0) at 37 °C for 20 min. After washing with PBS buffer (pH 7.4) and air-drying, soil samples were further treated with 10% (w/v) blocking reagent (Merck KGaA) in maleic acid buffer (100 mM maleic acid and 150 mM NaCl; pH 7.5) at 42 °C for 30 min. All samples were dehydrated using an ethanol series (50%, 80%, and 96% [v/v]) for 3 min each.

Hybridization was performed at 46 °C for 2 h using a buffer containing 0.9 M NaCl, 20 mM Tris-HCl (pH 8.0), 0.01% (w/v) SDS, formamide (0–70%, v/v), and probes (10 pmol/µL). Following hybridization, slides were immersed in washing buffers containing the corresponding formamide concentrations at 48 °C for 15 min. For soil samples, DNA was counterstained with 0.4 µg/mL 4′,6-diamidino-2-phenylindole (DAPI) (Wako) for 15 min. Finally, samples were mounted using ProLong Glass Antifade Mountant (Thermo Fisher Scientific).

### Microscopic observation, image analysis, and measurement of bacterial density

Observations were performed under oil immersion using a 100× objective lens on a BZ-9000 fluorescence microscope (Keyence, Osaka, Japan). The following filter sets were used: GFP-B (Keyence; excitation filter 470/40 nm, dichroic mirror 495 nm, and barrier filter 535/50 nm) for FITC; TRITC (Keyence; excitation filter 540/25 nm, dichroic mirror 565 nm, and barrier filter 605/55 nm) for ROX; and DAPI-5060C-KEY (Semrock; excitation filter 377/50 nm, dichroic mirror 409 nm, and barrier filter 447/60 nm) for DAPI. Images were processed using a BZ-II Analyzer version 1.42 (Keyence).

Images were analyzed using ImageJ Fiji version 2.1.0/1.53c (https://imagej.net/software/fiji/). For each bacterial cell, a circular region of interest (ROI) was defined to encompass the cell and its immediate background. The fluorescence intensity of the bacterial cells was determined by calculating the difference between the maximum and minimum intensities within the ROI. Bacterial density in the roots was calculated by dividing the cell count by the root surface area.

### Quantitative polymerase chain reaction (qPCR) analysis for measurement of bacterial density

Bacterial DNA was extracted from the root subsamples (250 mg per sample), which were prepared as described in “Sampling of roots and soil” using the DNeasy PowerSoil Pro Kit (QIAGEN, Hilden, Germany) according to the manufacturer’s instructions. qPCR was performed by targeting a portion of bacterial 16S rDNA. The universal primers (357F [5′-CCTACGGGAGGCAGCAG-3′] (Fierer *et al*. 2005) and 518R [5′-ATTACCGCGGCTGCTGG-3′] (Turner *et al*. 1999) were used to quantify the bacterial abundance of total bacteria. The abundance of *C. sedlakii* CESi7 was quantified using the *C. sedlakii*-specific primers (362F [5′-GGGTTGTGGTTAATAACCGCAGTC-3′] and 906R [5′- CATCTCTGCAGTCTTCCGTGG-3′]). Prior to qPCR analysis, the primer specificity was verified using PCR with 50 ng of genomic DNA from *C. sedlakii* CESi7 and *E. coli* DH5α as templates. PCR was performed using Ex Taq (Takara Bio, Shiga, Japan), according to the manufacturer’s instructions. The thermal cycling conditions consisted of an initial denaturation at 98 °C for 5 min, followed by 30 cycles of 98 °C for 30 s, 55 °C for 30 s, and 72 °C for 30 s, and a final elongation at 72 °C for 5 min. qPCR analysis was performed using each 10-μL reaction mixture consisting of 2 μL of KOD SYBR qPCR Mix (TOYOBO, Osaka, Japan), 0.1 μL of each primer, and 2 μL of bacterial DNA. The amplification program included an initial denaturation step at 95 °C for 5 min followed by 40 cycles of denaturation at 95 °C for 5 s, annealing at 64 °C for 20 s, and elongation at 72 °C for 20 s. DNA extracted from the strain CESi7 was used as the standard to generate a calibration curve. After amplification, the copy numbers of the 16S rDNA per root sample were calculated mm^−1^ of root and converted to mm^−2^ of root. The following equation was used for this calculation: Copies mm^-2^ of root = qPCR copies / [root length (mm) × root diameter (mm) × π]. Root diameter was measured using a stereoscope.

## RESULTS

### Design of probe targeting *C. sedlakii* CESi7

To achieve the in situ observation of *C. sedlakii* CESi7 in the roots and soil, we constructed a novel oligonucleotide probe targeting this strain. Five closely related type strains belonging to the genus *Citrobacter* and five type subspecies of *Salmonella enterica* were selected by a similarity search of the 16S rDNA sequences with the strain CESi7 (Table S1). After comparing alignments of their 16S rDNA sequences and confirming mismatched bases, the sequence binding to the region between positions 906 and 926 of the 16S rRNA was selected as a probe targeting strain CESi7, and it was named the CITR906 probe (Table 1). The designed CITR906 probe showed a 0.016% mishit rate with the following species of the respective type trains: 0 mer mismatch, four species; one mer mismatch, three species; two mer mismatches, 20 species (Table S2).

### Optimization of CITR906 hybridization conditions

The optimal formamide concentration for FISH with the CITR906 probe was determined using a pure culture of *C. sedlakii* CESi7. The fluorescence intensity of the CITR906 probe gradually increased as formamide concentration increased from 0 to 40%, followed by a decrease at higher concentrations (Fig. S1). Furthermore, the EUB338 probe, a universal probe for bacteria, exhibited a high affinity for CESi7 at 40% formamide (Fig. S1). Consequently, 40% formamide was used for subsequent FISH observations to detect CESi7 using both CITR906 and EUB338 probes.

To evaluate the specificity of the CITR906 probe, FISH observations were conducted at the optimized 40% formamide concentration using *R. tangshanensis*^T^, *M. theicola*^T^, and *E. coli* DH5α, which have one, two, and three mismatches to the CITR906 probe sequence, respectively (Table S2). These mismatches were located near the center of the probe sequence. The EUB338 probe provided sufficient fluorescence in all tested strains, indicating successful hybridization (Fig. S2). In contrast, the CITR906 probe exhibited barely detectable fluorescence for *M. theicola* and *E. coli* DH5α (Fig. S2). However, a distinct fluorescence was observed for strain CESi7 with the CITR906 probe, and *R. tangshanensis* cells were also labeled (Fig. S2), indicating that this probe can hybridize with 16S rRNA sequences containing up to a single-base mismatch. The potential impact of this probe’s hybridization with single-base-mismatched sequences is further addressed in the DISCUSSION section.

### FISH observation in *B. rapa* roots using sterilized medium

*B. rapa* was cultivated in a sterilized medium for one, three, and five weeks. FISH analysis revealed that *C. sedlakii* CESi7 colonized both the root tips and mature zones after one week of cultivation (Fig. 1). This strain was observed in the mature zones at week three and was absent in all root tissues by week five (Fig. S3). At one week of cultivation, strain CESi7 formed aggregated colonies throughout the mature zones, spanning the epidermis, root hairs, cortex, and vascular cylinder (Fig. 1A–D). Additionally, this strain was observed in the apical meristems of the root tips (Fig. 1E). The density of strain CESi7 calculated via FISH observations was significantly higher in the mature zones (0.9 × 10^4^ cells mm^-2^) than that in the root tips (0.6 × 10^4^ cells mm^-2^) (Welch’s *t*-test, *P* < 0.05; Table 2). No fluorescence signals from EUB338 or CITR906 were detected in the non-inoculated roots (Fig. 1F–I). These results demonstrate that strain CESi7 can colonize both the surface and internal tissues under aseptic conditions, exhibiting a significantly higher abundance in mature zones than in root tips.

**Fig. 1.**
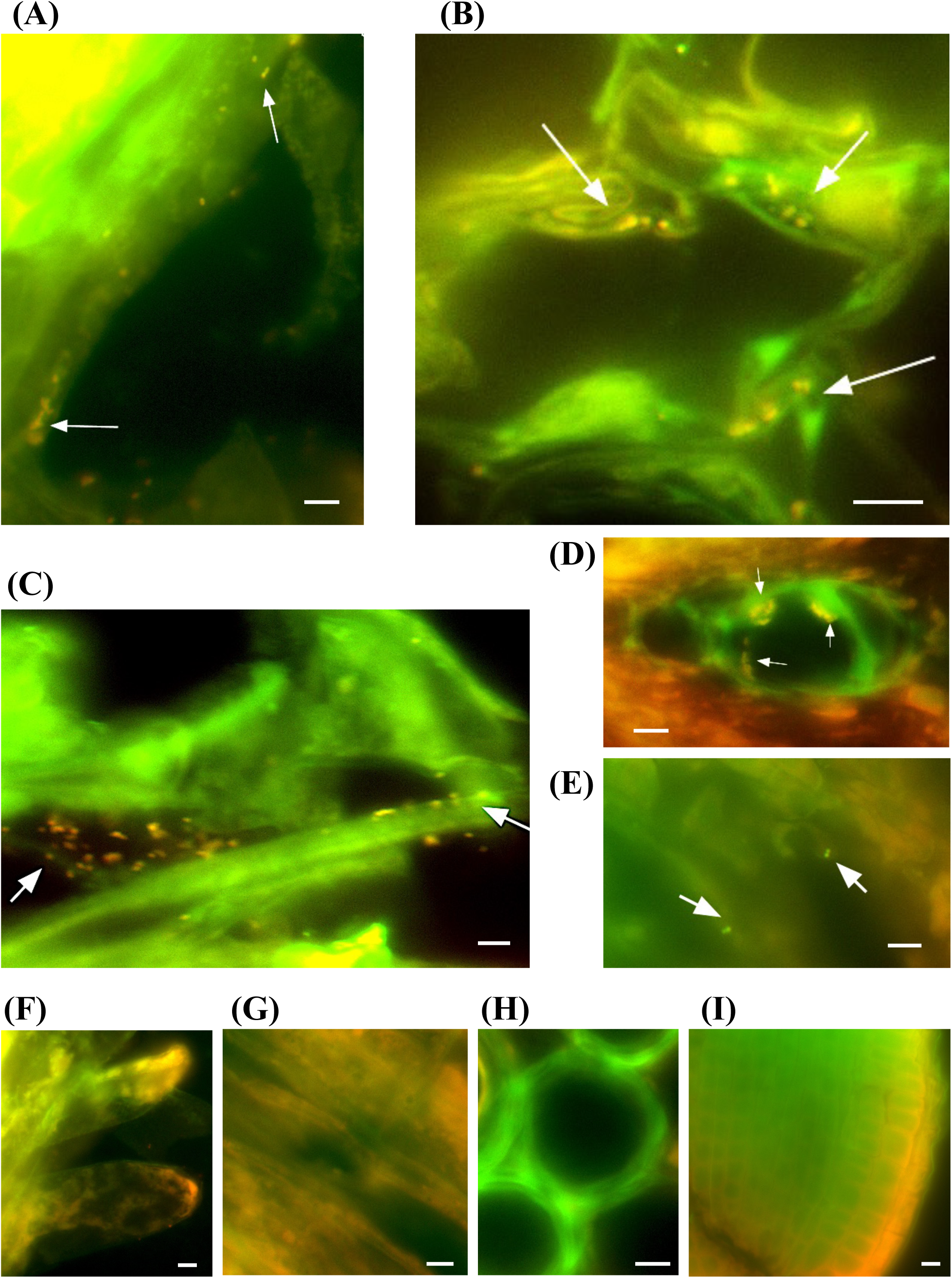
FISH analysis of *Brassica rapa* roots at one week cultivation after inoculation with or without *C. sedlakii* CESi7 in sterilized medium. (A–E) Root tissues inoculated with strain CESi7: (A) root hairs, (B) cortex, (C) epidermis, (D) vascular cylinder, and (E) root apical meristem. (F–I) Non-inoculated controls: (F) root hairs, (G) epidermis, (H) vascular cylinder, and (I) root apical meristem. All images are presented as overlays of green signals from the FITC-labelled CITR906 probe and red signals from the ROX-labelled EUB338 probe. Yellow-orange fluorescence indicates colonies of strain CESi7 (indicated by white arrows). Scale bars: 5 μm.

**Table 2.**
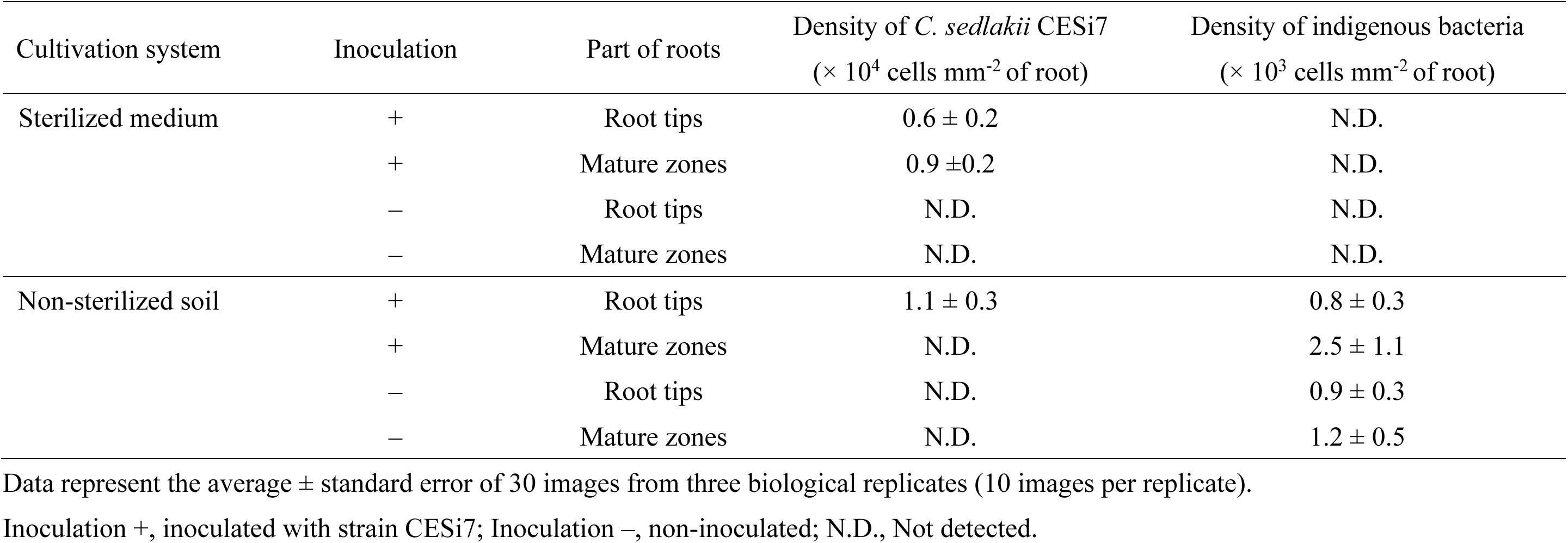
Bacterial density in *Brassica rapa* roots in one week of cultivation quantified by FISH observation.

| Cultivation system | Inoculation | Part of roots | Density of *C. sedlakii* CESi7<br>($\times 10^4$ cells mm<sup>-2</sup> of root) | Density of indigenous bacteria<br>($\times 10^3$ cells mm<sup>-2</sup> of root) |
| --- | --- | --- | --- | --- |
| Sterilized medium | + | Root tips | $0.6 \pm 0.2$ | N.D. |
| | + | Mature zones | $0.9 \pm 0.2$ | N.D. |
|  | – | Root tips | N.D. | N.D. |
|  | – | Mature zones | N.D. | N.D. |
| Non-sterilized soil | + | Root tips | $1.1 \pm 0.3$ | $0.8 \pm 0.3$ |
| | + | Mature zones | N.D. | $2.5 \pm 1.1$ |
| | – | Root tips | N.D. | $0.9 \pm 0.3$ |
| | – | Mature zones | N.D. | $1.2 \pm 0.5$ |

Data represent the average ± standard error of 30 images from three biological replicates (10 images per replicate).
Inoculation +, inoculated with strain CESi7; Inoculation –, non-inoculated; N.D., Not detected.

### FISH observation in *B. rapa* roots using non-sterilized soil

Bacterial colonization was monitored for five weeks during *B. rapa* cultivation in non-sterilized soil, both with and without inoculation with *C. sedlakii* CESi7. Following inoculation, CESi7 cells transiently colonized the root tips until the first week of cultivation (Fig. 2). Importantly, no yellow-orange fluorescence was observed in non-inoculated controls, confirming the absence of indigenous bacteria that cross-reacted with CITR906 (Fig. 2). Conversely, indigenous bacteria remained present across all sampled time points (one, three, and five weeks), regardless of inoculation with strain CESi7 (Fig. 2). Various root sections at one week of cultivation were subjected to FISH to compare the mature zones (Fig. 3AB) with the root tips (Fig. 3C–E). CESi7 formed several aggregated colonies within the root tips, including the root apical meristems (Fig. 3C), root caps (Fig. 3D), and the cortex within the root tips (Fig. 3E). Indigenous bacteria, except for strain CESi7, were observed in the epidermis (Fig. 3A) and root hairs (Figs. 3B and S4) of the mature zones. The density of strain CESi7 was 1.1 × 10^4^ cells mm^-2^ at the root tips in non-sterilized soil (Table 2). Furthermore, the densities of indigenous bacteria were comparable between the root tips and mature zones, averaging approximately 10^3^ cells mm^-2^, with no significant differences observed regardless of the inoculation of strain CESi7 (Welch’s *t*-test, *P* > 0.05; Table 2). These results indicate that strain CESi7 colonized both the surface and internal tissues of the root tips during the first week of cultivation in non-sterilized soil. Within the classification framework of colonization patterns, which includes symbiotic, endophytic, epiphytic, and free PGPB (Souza *et al*. 2015), our FISH observations demonstrate that strain CESi7 functions both as an endophytic and epiphytic PGPB colonizing the internal and external structures of *B. rapa* roots.

**Fig. 2.**
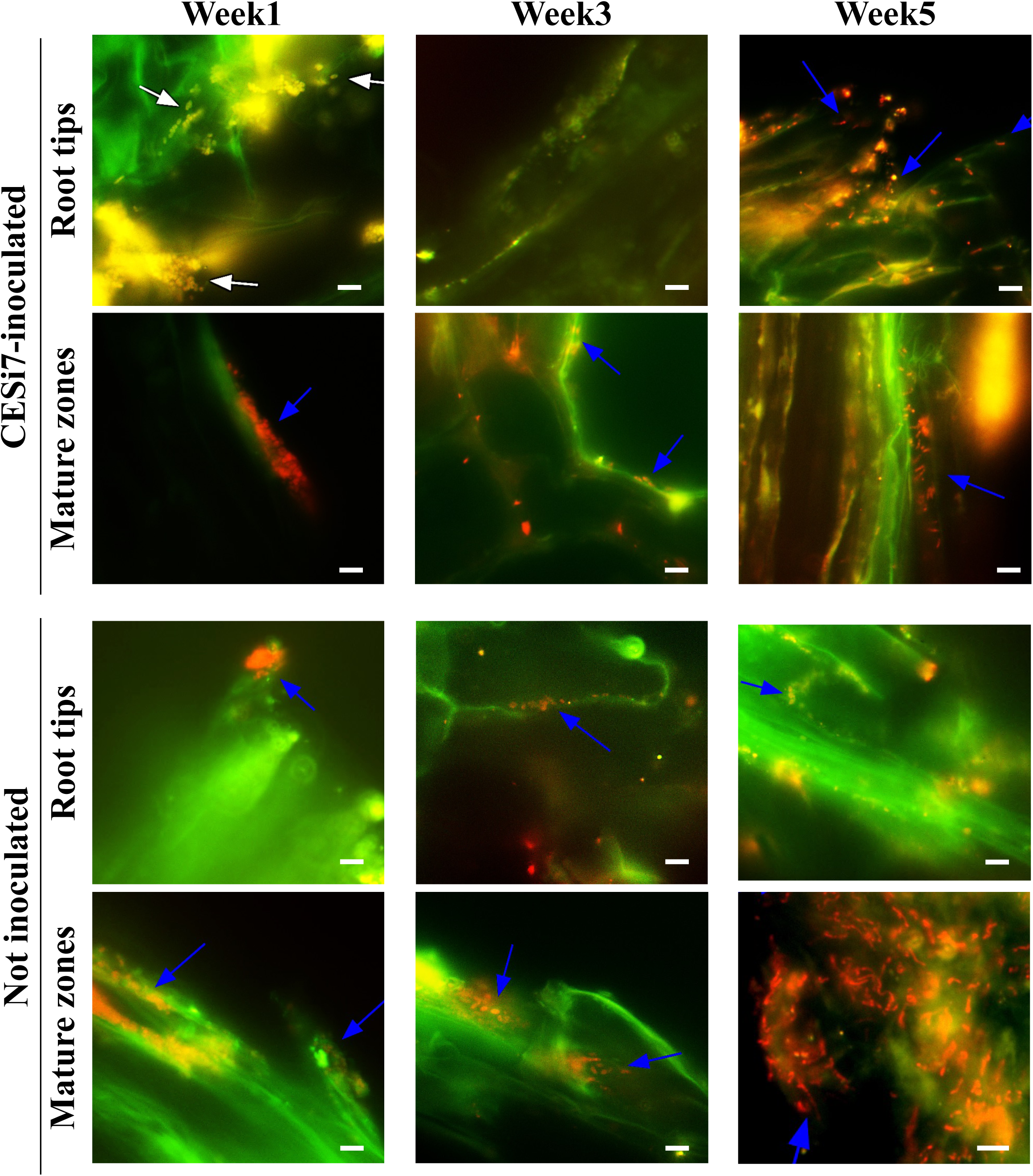
Spatiotemporal colonization of *C. sedlakii* CESi7 and indigenous bacteria in *Brassica rapa* roots after five weeks of cultivation in non-sterilized soil. All images are presented as overlays of green signals from the FITC-labelled CITR906 probe and red signals from the ROX-labelled EUB338 probe. Yellow-orange fluorescence indicates colonies of strain CESi7 (indicated by white arrows), while red fluorescence represents indigenous bacteria (indicated by blue arrows). Observations were conducted at both root tips and mature zones. Scale bars: 5 μm.

**Fig. 3.**
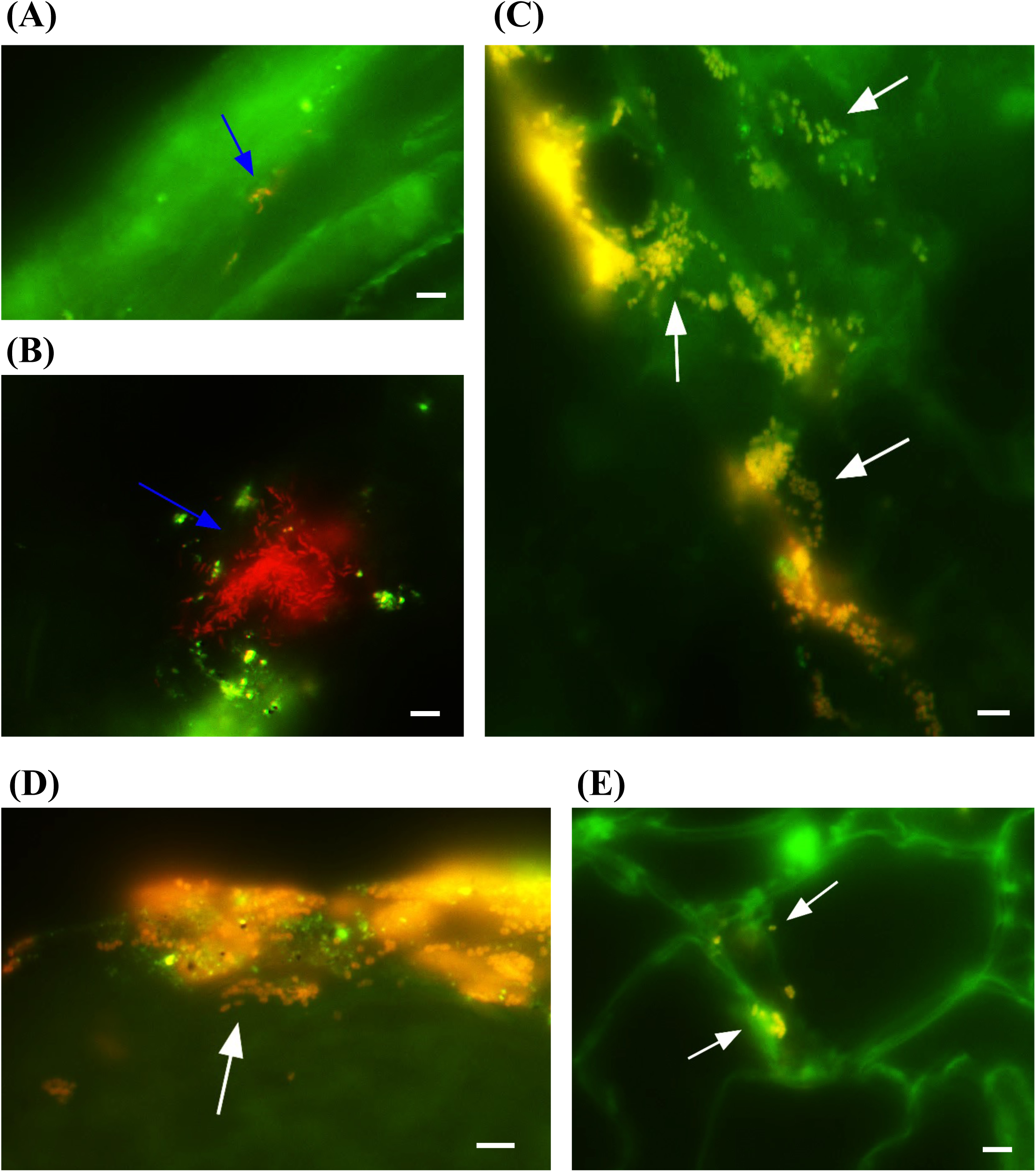
FISH observation of colonization patterns of *C. sedlakii* CESi7 and indigenous bacteria in *Brassica rapa* roots after one week cultivation with the inoculation in non-sterilized soil. (A–E) Representative images of different root tissues: (A) epidermis, (B) root hairs, (C) root apical meristem, (D) root cap, and (E) cortex. All images are presented as overlays of green signals from the FITC-labelled CITR906 probe and red signals from the ROX-labelled EUB338 probe. Yellow-orange fluorescence indicates colonies of strain CESi7 (indicated by white arrows), while red fluorescence represents indigenous bacteria (indicated by blue arrows). Scale bars: 5 μm.

### FISH observation in bulk and rhizosphere soils with *B. rapa* cultivation

To verify the visibility of CESi7 cells in the soil matrix, sterilized soil was mixed with a suspension of strain CESi7 and subjected to FISH analysis. The CESi7 cells were successfully visualized using the CITR906 probe in combination with DAPI staining (Fig. S5AC). To distinguish bacterial cells from autofluorescent abiotic soil particles, the negative control probe NONEUB, which binds non-specifically to soil particles but not to bacterial cells (Fig. S5B), was utilized. In the merged images of FITC-labeled CITR906, DAPI, and ROX-labeled NONEUB, the CESi7 cells were identified based on their cyan appearance (Fig. S5D). This approach allowed the clear visualization of strain CESi7 within the soil matrix.

The colonization of *C. sedlakii* CESi7 in non-sterilized soil was examined. However, no cyan bacterial cells, resulting from the overlay of FITC-labeled CITR906 (green) and DAPI (blue) fluorescence, were observed in either rhizosphere (Fig. S6) or bulk (Fig. S7) soil after one, three, or five weeks of cultivation. Indigenous microorganisms were consistently detected throughout the cultivation period (Figs. S6 and S7). These results indicate that strain CESi7 does not form detectable colonies within the soil during *B. rapa* cultivation.

### Quantification of bacterial abundance in *B. rapa* roots via qPCR analysis

The specificity of the primers was validated via PCR using genomic DNA from *C. sedlakii* CESi7 and *E. coli* DH5α as templates. The universal primers 357F and 518R yielded PCR products of the expected size (ca. 160 bp) for both strains (Fig. S8A). The CESi7-targeted primers, 362F and 906R, produced a target-sized fragment (ca.540 bp) with the strain CESi7 template, and no amplification was observed with the strain DH5α template (Fig. S8B). Therefore, the universal and CESi7-targeting primers were used for subsequent qPCR analyses of the total bacteria and strain CESi7, respectively.

To quantify bacterial abundance, qPCR targeting 16S rDNA was performed on DNA extracted from root samples after one week of cultivation (Tables S3 and S4). Under sterilized conditions, strain CESi7 was detected in both the root tips and mature zones with densities of 1.7 × 10^5^ and 0.4 × 10^5^ copies mm^−2^, respectively (Table S3). The abundance of total bacteria in the CESi7-inoculated samples was greater than that in the strain CESi7, and bacterial signals were detected in the non-inoculated control samples (Table S3). The quantified density of strain CESi7 was 5.9 copies mm^−2^ in the root tips and 0.8 copies mm^−2^ in the mature zones under non-sterilized conditions (Table S4).

## DISCUSSION

PGPB are increasingly recognized for their potential as biofertilizers, contributing to more sustainable agricultural practices by reducing the reliance on chemical fertilizers (Bashan *et al*. 1998; Gupta *et al*. 2015). Our previously isolated strain, *C. sedlakii* CESi7, was shown to enhance the growth of *B. rapa* (Chin *et al*. 2017); however, its colonization dynamics within roots and soil remained elusive. In this study, we designed a FISH probe targeting the strain CESi7 and demonstrated that this strain forms an epiphytic and endophytic population in the roots of *B. rapa*. Four species within the genus *Citrobacter*, other than *C. sedlakii* have been reported as PGPB: *Citrobacter freundii* (Nguyen *et al*. 2003), *Citrobacter youngae* (Nongkhlaw *et al*. 2014), *Citrobacter murliniae* (Kisiel *et al*. 2016), and *Citrobacter braakii* (Denaya *et al*. 2021). To the best of our knowledge, this is the first study to elucidate the colonization patterns of PGPB belonging to the genus *Citrobacter* during plant cultivation using direct microscopic observations.

The designed CITR906 probe could hybridize with bacteria possessing a single-base mismatch in their target sequences (Fig. S2). This potential binding includes several species, particularly *C. youngae*^T^, *Shewanella oneidensis*^T^, *Nitrincola alkalisediminis*^T^, *Rheinheimera soli*^T^, *Rheinheimera mesophila*^T^, and *R. tangshanensis*^T^ (Table S2). Among these, *C. youngae*^T^ and *R. soli*^T^ have been reported to inhabit soil environments (Ryu *et al*. 2008; Nongkhlaw *et al*. 2014), suggesting that their potential detection as false positives could not be entirely ruled out during observations in non-sterilized soil. However, no yellow-orange fluorescence was observed in the non-inoculated samples throughout the five-week cultivation period (Fig. 2), confirming that indigenous bacteria did not interfere with the specific detection of strain CESi7 in this study.

In addition to the FISH observations, bacterial abundance was quantified using qPCR analysis (Tables S3 and S4). In sterilized medium cultivation, the observed discrepancy between total and CESi7-specific counts, coupled with signals detected in non-inoculated controls (Table S3), could be attributed to two factors. First, universal 16S rRNA gene primers may cross-react with plant organellar DNA such as mitochondrial or chloroplast sequences (Edwards *et al*. 2007; Giangacomo et al. 2021). Second, the residual bacterial DNA derived from dead cells was amplified even after seed surface sterilization. These background signals likely led to an overestimation of the total bacterial abundance. During non-sterilized soil cultivation, the density of strain CESi7 within the *B. rapa* root samples quantified via qPCR (0.8–5.9 copies mm^-2^) was markedly lower than that obtained using FISH observation (1.1 × 10^4^ cells mm^-2^) (Tables 2 and S4). Although FISH clearly visualizes CESi7 colonization, the difficulty of recovering high-quality bacterial DNA from the complex root matrix (Giangacomo *et al*. 2021) may have led to an underestimation of qPCR values. Therefore, image-based FISH analysis more accurately reflects the actual colonization density of strain CESi7, with the additional advantage of identifying specific colonization sites in the root tissue.

Information on the time-scale colonization pattern and bacterial density of strain CESi7 could reveal the mechanisms through which it promotes plant growth. Strain CESi7 was observed at a density of 1.1 × 10^4^ cells mm^-2^ at the root tips only after the first week of cultivation using non-sterilized soil, after which its population diminished (Table 2 and Fig. 2). This suggests that strain CESi7 plays a key role in plant growth promotion at the early stage of *B. rapa* cultivation. Strain CESi7 was no longer detectable by week five, even in sterilized medium (Fig. S3), suggesting that the transient nature of colonization is an inherent characteristic of this strain, rather than a result of competitive exclusion by other bacteria.

The present FISH observations indicated the ability of strain CESi7 to colonize both the internal and external root tissues of *B. rapa*, characterizing it as an endophytic and epiphytic PGPB (Figs. 1 and 3). Several factors contribute to the early colonization of endophytic PGPB at specific sites, such as the crack entry process via root formation sites and the root tip pathway via root tips (Reinhold-Hurek *et al*. 1998; Compant *et al*. 2010). Strain CESi7 invaded the internal tissues of the root through the crack entry process and root tip pathway under sterilized medium cultivation, whereas the only invasion pathway under non-sterilized soil cultivation was the root tip pathway. The factors that cause these differences in the colonization sites of strain CESi7, depending on the cultivation system, warrant systematic investigation in future studies.

In addition to strain CESi7, the colonization patterns of PGPB belonging to *Enterobacteriaceae*, including *Enterobacter* (Gontia-Mishra *et al*. 2016; Subrahmanyam *et al*. 2018; Costa Júnior *et al*. 2020; Galambos *et al*. 2020), *Klebsiella* (Gontia-Mishra *et al*. 2016; Sapre *et al*. 2018), *Kosakonia* (Witzel *et al*. 2017, Singh *et al*. 2020), and *Raoultella* (Luo *et al*. 2016), have been elucidated (Table 3). Almost all PGPB except *Enterobacter* sp. C1D (Subrahmanyam *et al*. 2018) and *Kosakonia radicincitrans* DSM 16656^T^ (Witzel *et al*. 2017) colonize the interior of roots as endophytes. Because endophytic PGPB exhibit greater plant growth-promoting activity than their epiphytic counterparts (Chanway *et al*. 2000), many strains within the *Enterobacteriaceae* family are considered promising candidates for biofertilizer development. However, this study elucidates the actual colonization dynamics of an *Enterobacteriaceae* PGPB strain with non-sterilized soil in the presence of indigenous bacteria (Table 3).

**Table 3.** Colonization pattern of PGPB family *Enterobacteriaceae*.

| PGPB | Host plant | Observation method | Cultivation system | Colonization sites | Plants/bacteria interaction types | Reference |
| --- | --- | --- | --- | --- | --- | --- |
| *Citrobacter sedlakii* CESi7 | *Brassica rapa* | FISH-Fluorescence microscope | Non-sterilized soil;<br>Sterilized medium | (Non-sterilized soil)<br>Root surface and intercellular space of the root tip zones;<br>(Sterilized medium)<br>Root surface and interior | Endophytic, epiphytic | This study |
| *Enterobacter cloacae* M18B;<br>*Enterobacter cloacae* M19B;<br>*Enterobacter cloacae* CCMA1285 | Garlic | SEM | Sterilized medium | Surface and interior of the roots | Endophytic, epiphytic | (Costa Júnior *et al.* 2020) |
| *Enterobacter* sp. 32A | Tomato | FISH-CLSM | Sterilized medium | Surface and interior of the lateral root formation sites,<br>root tips, elongation zones; Root hairs; Xylems | Endophytic, epiphytic | (Galambos *et al.* 2020) |
| *Enterobacter* sp. C1D | Mung bean | GFP label-CLSM | Sterilized medium | Root surface of the root tip zones;<br>Root hair/lateral root formation sites | Epiphytic | (Subrahmanyam *et al.* 2018) |
| *Enterobacter ludwigii* IG10 | Wheat | SEM; TTC staining* | Sterilized soil | Surface and interior of the roots | Endophytic, epiphytic | (Gontia-Mishra *et al.* 2016) |
| *Klebsiella* sp. IG3 | Wheat | SEM; TTC staining* | Sterilized soil | Surface and interior of the roots | Endophytic, epiphytic | (Gontia-Mishra *et al.* 2016) |
| *Klebsiella* sp. IG3 | Oat | SEM; TTC staining* | Sterilized medium | Surface and interior of the roots | Endophytic, epiphytic | (Sapre *et al.* 2018) |
| *Kosakonia radicincitans* DSM 16656<sup>T</sup> | *Arabidopsis thaliana* | GFP label-CLSM | Sterilized medium | Root surface; Root hair; Lateral root formation sites | Epiphytic | (Witzel *et al.* 2017) |
| *Kosakonia radicincitans* BA1 | Sugarcane | GFP label-CLSM; SEM | Sterilized medium | Surface and interior of the roots | Endophytic, epiphytic | (Singh *et al.* 2020) |
| *Raoultella* sp. L03 | Sugarcane | GFP label-Fluorescence microscope | Sterilized medium | Cortical intercellular spaces at the root maturation zone | Endophytic, epiphytic | (Luo *et al.* 2016) |

TTC staining\*: 2, 3, 5-triphenyl tetrazolium chloride staining

In conclusion, this study visualized the colonization patterns of the novel PGPB strain *C. sedlakii* CESi7, demonstrating its dual nature as both an endophytic and epiphytic bacterium. These findings provide a critical basis for optimizing the inoculation timing and application strategies to maximize its efficacy as a biofertilizer.

## Author contribution

Hiromu Inoue, Conceptualization, Investigation, Writing – original draft | Manami Maeda, Investigation | Tomonori Koga, Investigation | Zahra Salman, Investigation | Clament Fui Seung Chin, Resources | Mohd. Huzairi Mohd. Zainudin, Resources | Norhayati Bt. Ramli, Resources | Mohd Ali Hassan, Resources | Yukihiro Tashiro, Conceptualization, Project administration, Resources, Writing – review and editing | Kenji Sakai, Conceptualization, Project administration, Resources, Writing – review and editing

## Acknowledgments

The authors would like to acknowledge the Center for advanced instrumental and educational supports (Faculty of Agriculture, Kyushu University, Fukuoka, Japan) for the cryostat to prepare the frozen section, and the Environmental Control Center for Experimental Biology (Kyushu University, Fukuoka, Japan) for the provision of greenhouse.

