## Supplemental Figs 1-8, Tables 1-4 for "Fluorescence *in situ* hybridization reveals endophytic and epiphytic root colonization of the novel plant growth-promoting bacterium *Citrobacter sedlakii* CESi7"

#### Supporting figures

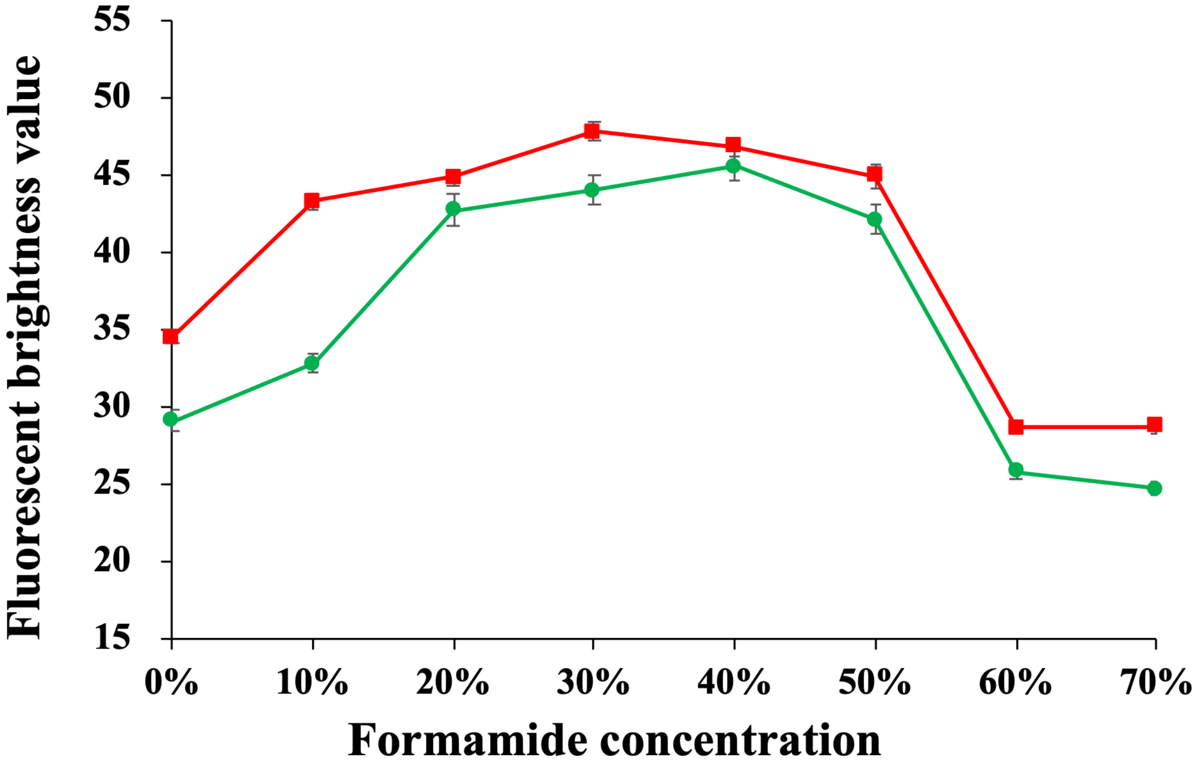


Fig. S1 Change of the fluorescent brightness value from the CITR906 probe and the EUB338 probe under *C. sedlakii* CESi7 pure cultures with several formamide concentrations. Symbols represent average value of fluorescent intensity form fifty cells of triplicate ($n$ = 150). Avg +/- SE. Symbols: closed circle, CITR906; closed square, EUB338.

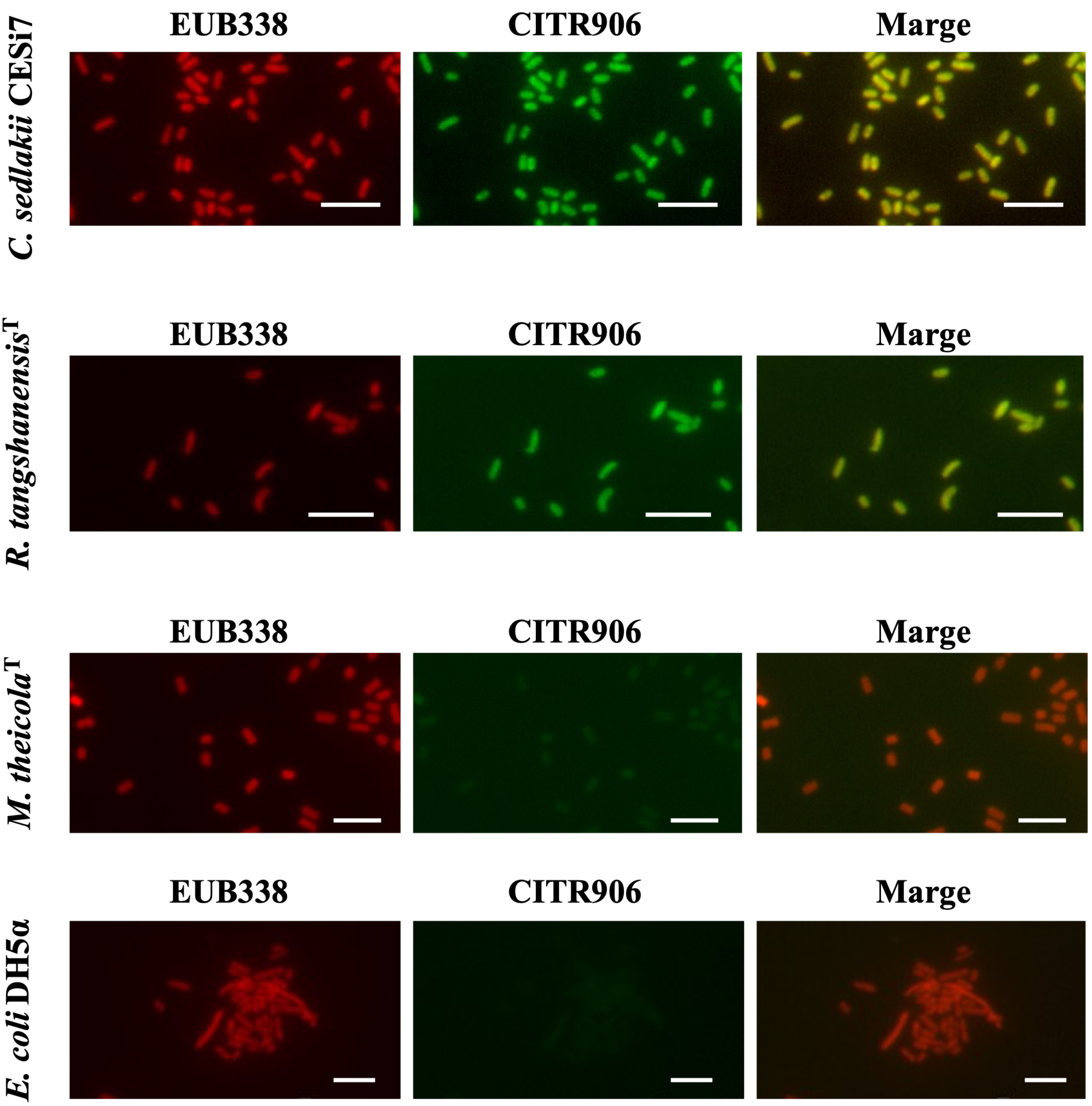


Fig. S2 FISH analysis of *C. sedlakii* CESi7, *R. tangshanensis*$^{T}$, *M. theicola*$^{T}$, and *E. coli* DH5α in pure cultures. Hybridization was performed using the EUB338 and CITR906 probes. All microphotographs were captured using a constant exposure time of 1 s to allow for comparison of fluorescence intensities. Scale bars: 5 μm.

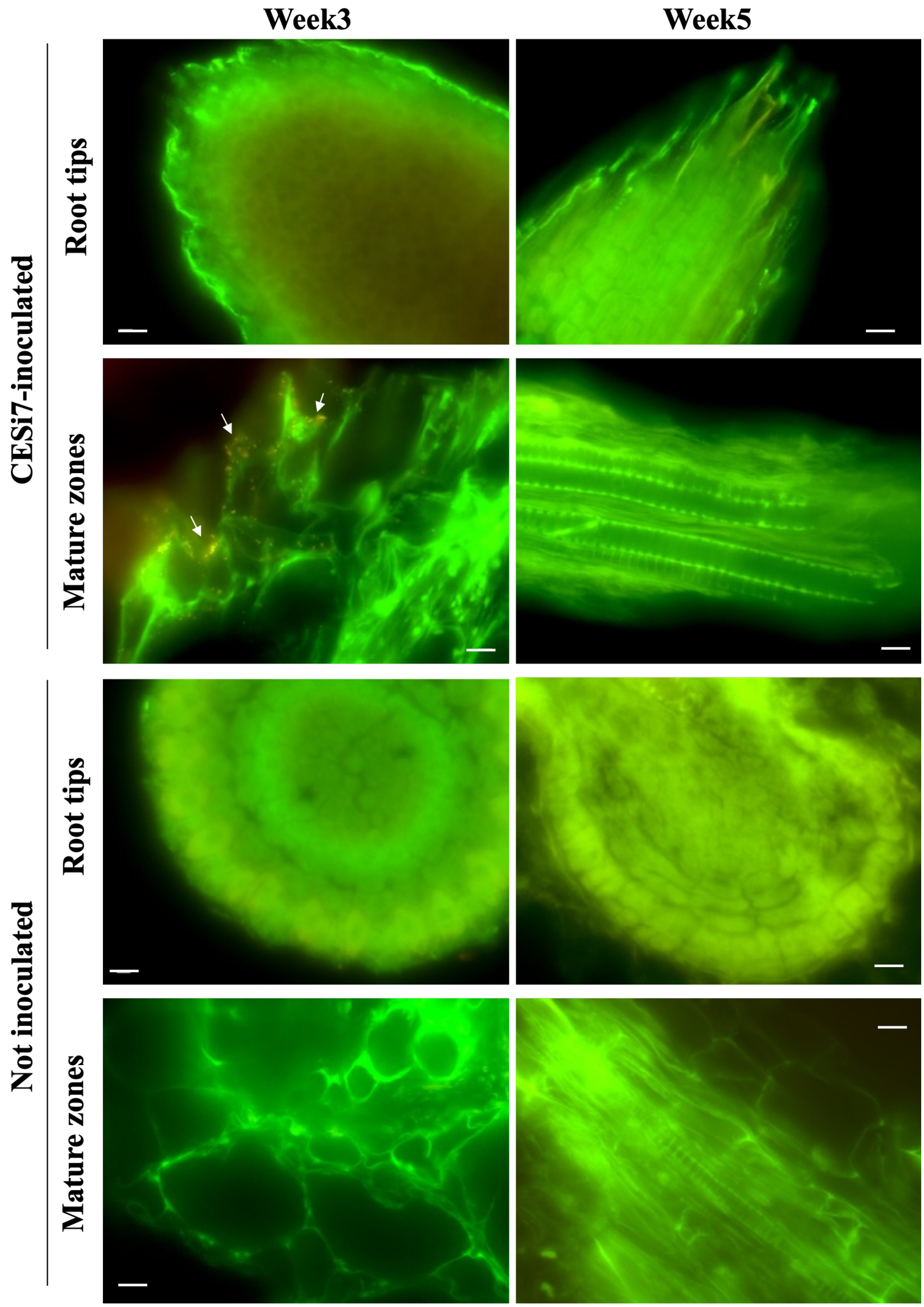


Fig. S3 Spatiotemporal colonization of *C. sedlakii* CESi7 on root tips and mature zones over five weeks of cultivation in sterilized medium. All images are presented as overlays of green signals from the FITC-labelled CITR906 probe and red signals from the ROX-labelled EUB338 probe. Yellow-orange fluorescence indicates the colonies of strain CESi7 (highlighted by white arrows). Scale bars: 10 μm.

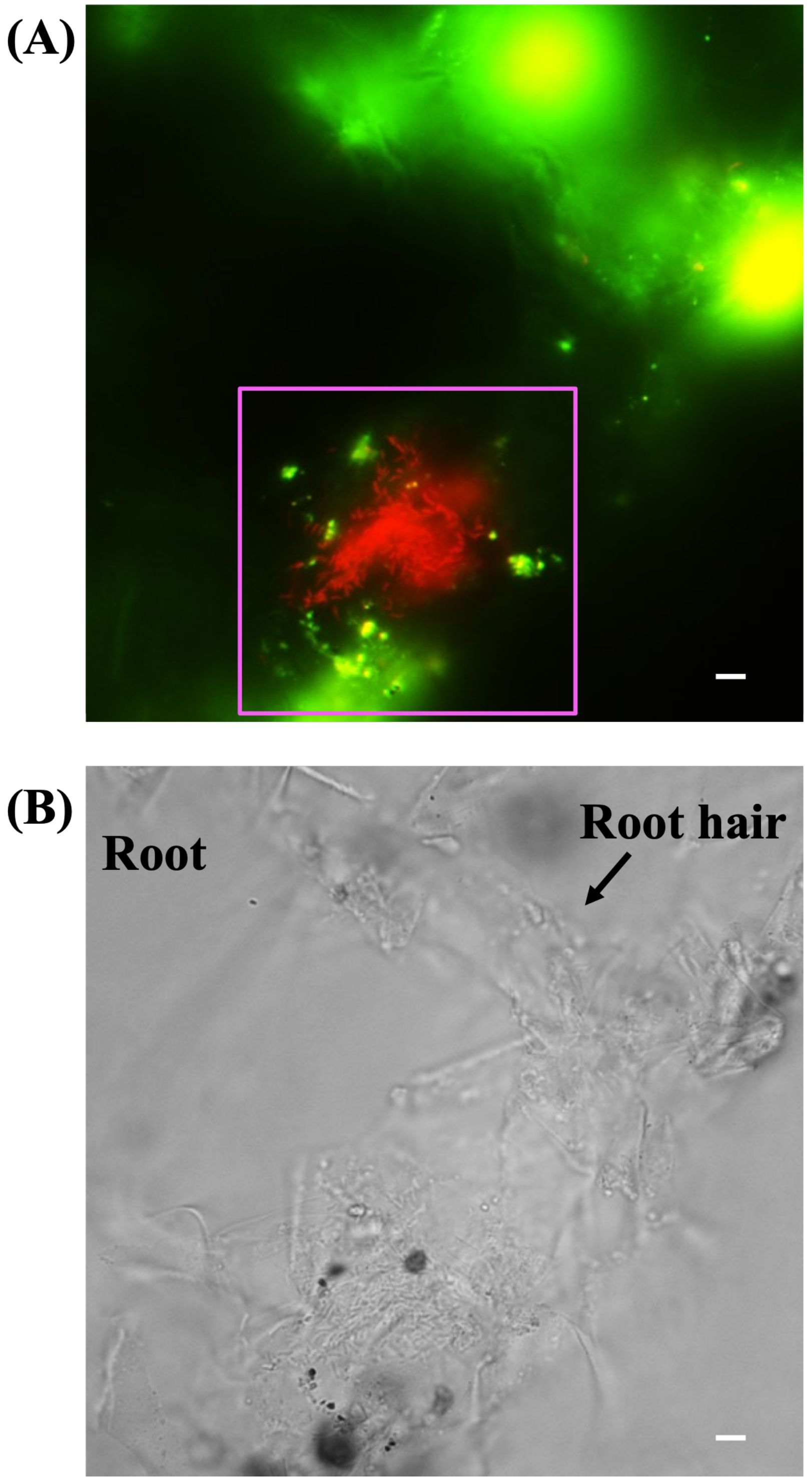


Fig. S4 Representative uncropped FISH and bright-field images of the root hairs shown in Fig. 3B. (A) Double-staining FISH image presented as an overlay of green signals from the FITC-labelled CITR906 probe and red signals from the ROX-labelled EUB338 probe. The magenta square indicates the specific region cropped and featured in Fig. 3B. (B) Corresponding bright-field image of the same field of view. Scale bars: 5 µm.

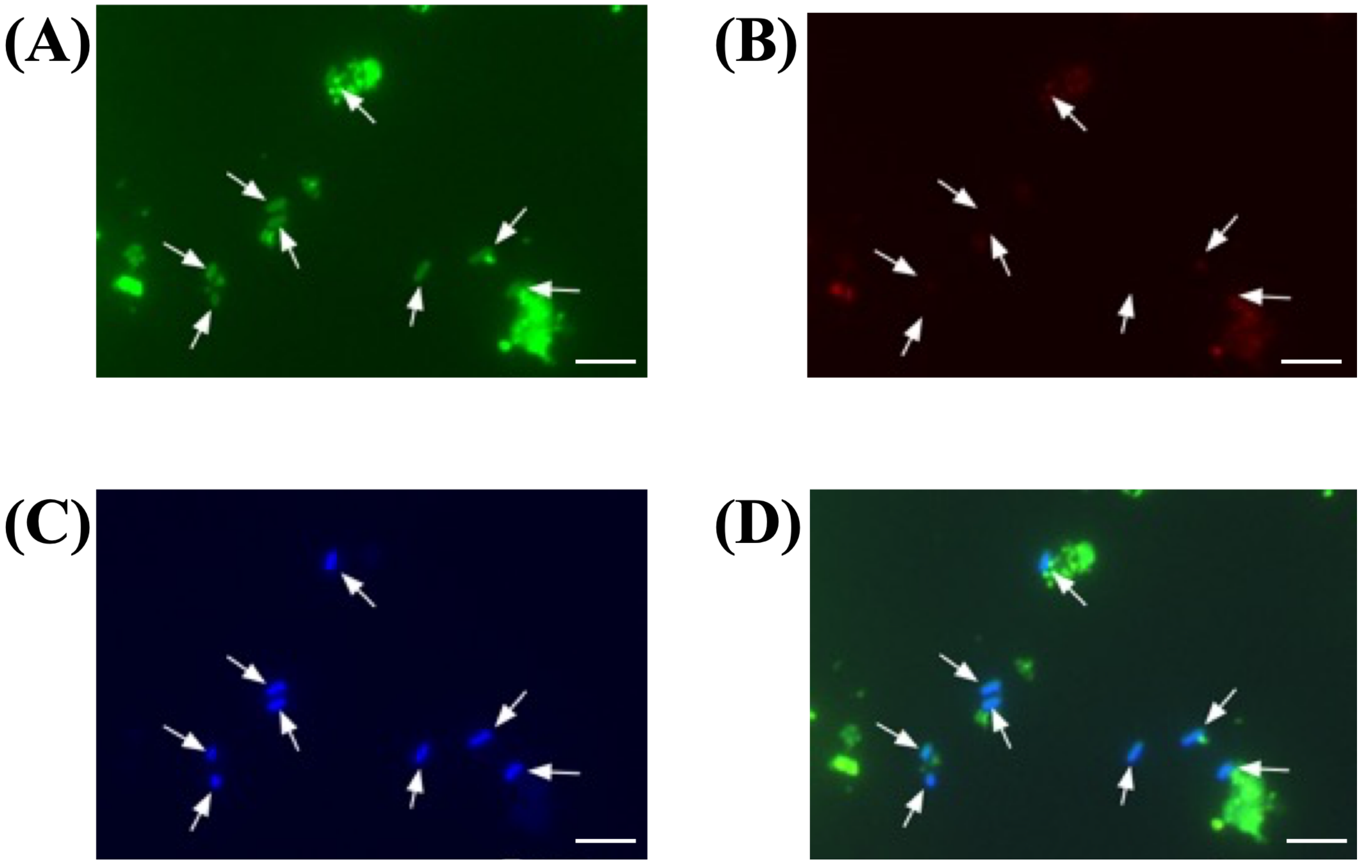


Fig. S5 FISH analysis of sterilized soil supplemented with *C. sedlakii* CESi7 suspension. (A) Green signals from the FITC-labelled CITR906 probe. (B) Red signals from the ROX-labelled NONEUB probe (negative control). (C) Blue signals from DAPI staining (total DNA). (D) Overlaid image of panels (A), (B), and (C). White arrows indicate the cells of strain CESi7, which appear cyan in the overlay due to the co-localization of FITC and DAPI signals. Scale bars: 5 µm.

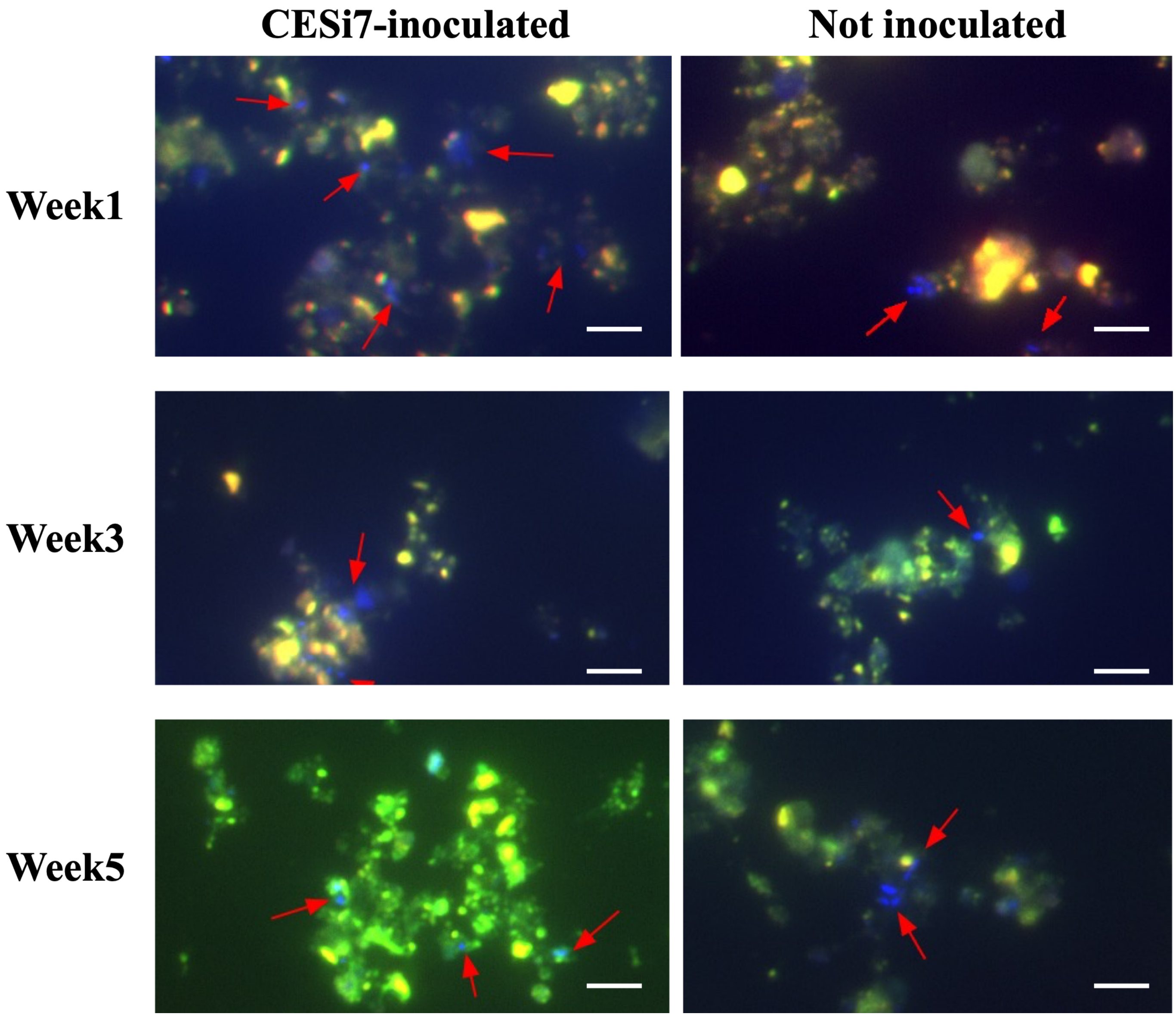


Fig. S6 FISH analysis of rhizosphere soil at 1, 3, and 5 weeks of cultivation in non-sterilized soil. All images are presented as overlays of green signals from the FITC-labelled CITR906 probe, red signals from the ROX-labelled NONEUB probe (negative control), and blue signals from DAPI staining (total DNA). Blue-labelled cells (indicated by red arrows) represent indigenous microorganisms, which were stained only by DAPI. Scale bars: 5 μm.

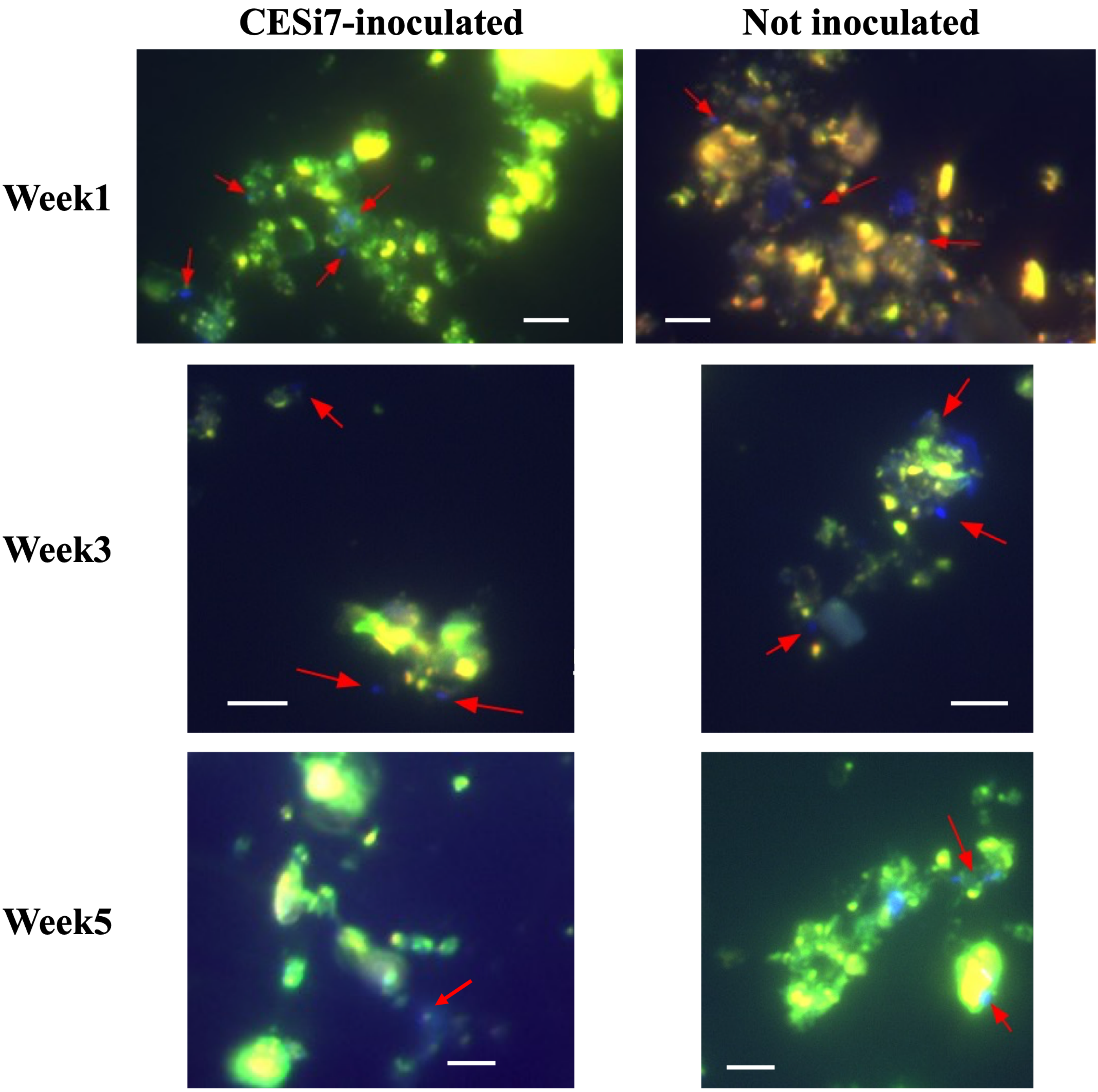


Fig. S7 FISH analysis of bulk soil at 1, 3, and 5 weeks of cultivation in non-sterilized soil. All images are presented as overlays of green signals from the FITC-labelled CITR906 probe, red signals from the ROX-labelled NONEUB probe (negative control), and blue signals from DAPI staining (total DNA). Blue-labelled cells (indicated by red arrows) represent indigenous microorganisms, which were stained only by DAPI. Scale bars: 5 μm.

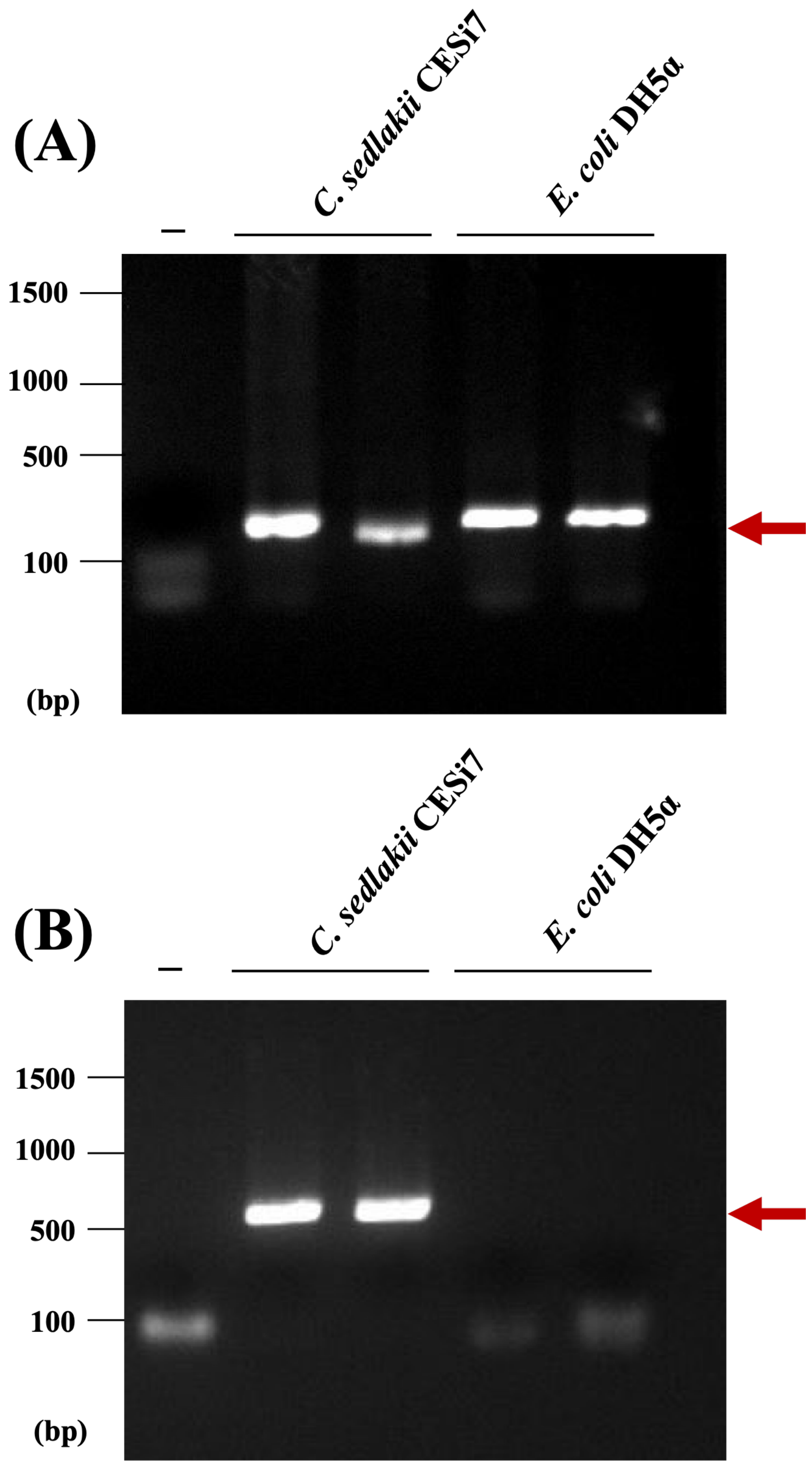


Fig. S8 Verification of primer specificity for qPCR analysis. Agarose gel electrophoresis of PCR products amplified using (A) universal primers, 357F and 518R, and (B) CESi7-targeted primers, 362F and 906R. Red arrows indicate the expected size of the respective PCR products. The lane labelled "–" represents the negative control (no-template control). The experiment was performed in biological duplicate.

### Supporting tables

Table S1 16S rDNA sequence similarities of *C. sedlakii* CESi7 with its closely related type strains

| Rank | Species | Pairwise similarity (%) | Mismatch / Total nt |
|---|---|---|---|
| 1 | *Citrobacter sedlakii*$^{T}$ | 99.70 | 4/1342 |
| 2 | *Citrobacter rodentium*$^{T}$ | 98.73 | 17/1342 |
| 3 | *Citrobacter farmeri*$^{T}$ | 98.58 | 19/1341 |
| 4 | *Citrobacter amalonaticus*$^{T}$ | 98.36 | 22/1342 |
| 5 | *Salmonella enterica* subsp. *salamae*$^{T}$ | 98.36 | 22/1341 |
| 6 | *Salmonella enterica* subsp. *diarizonae*$^{T}$ | 98.29 | 23/1342 |
| 7 | *Citrobacter koseri*$^{T}$ | 98.28 | 23/1336 |
| 8 | *Salmonella enterica* subsp. *indica*$^{T}$ | 98.14 | 25/1341 |
| 9 | *Salmonella enterica* subsp. *houtenae*$^{T}$ | 97.99 | 27/1341 |
| 10 | *Salmonella enterica* subsp. *arizonae*$^{T}$ | 97.76 | 30/1341 |

Table S2 Sequence mismatches of the CITR906 probe against the 16S rDNA of selected type strains

| The number of mismatches | Hit/All type strain | Type strain name |
|---|---|---|
| 0 | 4/12589 | *Citrobacter sedlakii*$^{T}$, *Citrobacter youngae*$^{T}$, *Shewanella oneidensis*$^{T}$, *Nitrincola alkalisediminis*$^{T}$ |
| 1 | 3/12589 | *Rheinheimera soli*$^{T}$, *Rheinheimera mesophila*$^{T}$, *Rheinheimera tangshanensis*$^{T}$ |
| 2 | 20/12589 | *Erwinia amylovora*$^{T}$, *Erwinia mallotivora*$^{T}$, *Dickeya paradisiaca*$^{T}$, *Erwinia papayae*$^{T}$, *Microbulbifer variabilis*$^{T}$, *Photobacterium ganghwense*$^{T}$, *Xenorhabdus indica*$^{T}$, *Rheinheimera texasensis*$^{T}$, *Microbulbifer epialgicus*$^{T}$, *Xenorhabdus griffiniae*$^{T}$, *Shewanella algicola*$^{T}$, *Rheinheimera aquatica*$^{T}$, *Erwinia piriflorinigrans*$^{T}$, *Halomonas daqiaonensis*$^{T}$, *Erwinia uzenensis*$^{T}$, *Xenorhabdus ishibashii*$^{T}$, *Photobacterium ganghwense*$^{T}$, *Microbulbifer echini*$^{T}$, *Mixta theicola*$^{T}$, *Nautilia abyssi*$^{T}$ |

Table S3

Bacterial density in sterilized medium for one week *Brassica rapa* cultivation quantified by qPCR analysis

| Inoculation | Part of roots | Density of *C. sedlakii* CESi7 ($\times 10^5$ copies $mm^{-2}$ of root) | Density of total bacteria ($\times 10^5$ copies $mm^{-2}$ of root) |
|---|---|---|---|
| + | Root tips | 1.7 ± 1.3 | 6.2 ± 2.5 |
| + | Mature zones | 0.4 ± 0.2 | 0.7 ± 0.4 |
| – | Root tips | N.D. | 0.3 ± 0.2 |
| – | Mature zones | N.D. | 1.1 ± 0.7 |

Data represent the average ± standard error of three biological replicates.

Inoculation +, inoculated with strain CESi7; Inoculation –, non-inoculated; N.D., Not detected.

Table S4

Bacterial density in non-sterilized soil for one week *Brassica rapa* cultivation quantified by qPCR analysis

| Inoculation | Part of roots | Density of *C. sedlakii* CESi7 (copies $mm^{-2}$ of root) | Density of total bacteria ($\times 10^2$ copies $mm^{-2}$ of root) |
|---|---|---|---|
| + | Root tips | 5.9 ± 4.8 | 8.3 ± 5.3 |
| + | Mature zones | 0.8 ± 0.6 | 0.7 ± 0.3 |
| – | Root tips | N.D. | 45.0 ± 30.0 |
| – | Mature zones | N.D. | 1.5 ± 1.0 |

Data represent the average ± standard error of three biological replicates.

Inoculation +, inoculated with strain CESi7; Inoculation –, non-inoculated; N.D., Not detected.
